# Local chromosome context is a major determinant of crossover pathway biochemistry during budding yeast meiosis

**DOI:** 10.1101/064519

**Authors:** Darpan Medhi, Alastair S. H. Goldman, Michael Lichten

## Abstract

Meiotic chromosomes are divided into regions of enrichment and depletion for meiotic chromosome axis proteins, in budding yeast Hop1 and Red1. These proteins are important for formation of Spo11-catalyzed DSB, but their contribution to crossover recombination is undefined. By studying meiotic recombination initiated by the sequence-specific *VMA1*-derived endonuclease (VDE), we show that meiotic chromosome structure helps to determine the biochemical mechanism by which recombination intermediates are resolved to form crossovers. At a Hop1-enriched locus, most VDE-initiated crossovers required the MutLγ resolvase, which forms most Spo11-initiated crossovers. In contrast, at a locus with lower Hop1 occupancy, most VDE-initiated crossovers were MutLγ-independent. In *pch2* mutants, the two loci displayed similar Hop1 occupancy levels, and also displayed similar MutLγ-dependence of VDE-induced crossovers. We suggest that meiotic and mitotic recombination pathways coexist within meiotic cells, with features of meiotic chromosome structure partitioning the genome into regions where one pathway or the other predominates.

## Introduction

The transition from the mitotic cell cycle to meiosis involves substantial changes in mechanisms of DNA double strand break (DSB) repair by homologous recombination (HR). Most mitotic HR repairs spontaneous lesions, and most repair products are non-crossovers (NCOs) that do not involve exchange of flanking parental sequences (Fabre et al., 1984; Kadyk and Hartwell, 1992) (Ira et al., 2003; Kelly, 1974; Pâques et al., 1998; Stark and Jasin, 2003; Taghian and Nickoloff, 1997; Virgin et al., 2001). In contrast, meiotic recombination is initiated by programmed DSBs (Cao et al., 1990; Sun et al., 1989) that often are repaired as crossovers (COs) between homologous chromosomes (homologs), with exchange of flanking parental sequences. Inter-homolog COs create physical linkages, called chiasmata, that ensure faithful homolog segregation during the first meiotic division, avoiding chromosome nondisjunction and consequent aneuploidy in gametes [reviewed by (Hunter, 2015)].

The DSBs that initiate meiotic recombination are formed by Spo11 in complex with a number of accessory proteins, and will be referred to here as Spo11-DSBs [reviewed by (Lam and Keeney, 2015)]. Spo11-DSBs and resulting recombination events are non-uniformly distributed in the genomes of organisms ranging from budding yeast to humans (Baudat Nicolas, 1997; Blitzblau et al., 2007; Buhler et al., 2007; Gerton et al., 2000; Pan et al., 2011) (Fowler et al., 2013) (Wijnker et al., 2013) (Hellsten et al., 2013) (Singhal et al., 2015) (Smagulova et al., 2011) (Pratto et al., 2014). In budding yeast, this nonuniform distribution of Spo11-DSBs is influenced by meiosis-specific proteins, Red1 and Hop1, which are components of the meiotic chromosome axis. The meiotic chromosome axis coordinates sister chromatids and forms the axial element of the synaptonemal complex, which holds homologs in tight juxtaposition (Hollingsworth et al., 1990; Page and Hawley, 2004; Smith and Roeder, 1997; Sym et al., 1993). Spo11-DSBs form frequently in large (ca 50-200 kb) “hot” domains that are also enriched for Red1 and Hop1, and these “hot” domains are interspersed with similarly-sized “cold” regions where Spo11-DSBs are infrequent and Red1/Hop1 occupancy levels are low (Baudat and Nicolas, 1997; Blat et al., 2002; Blitzblau et al., 2007; Buhler et al., 2007; Pan et al., 2011; Panizza et al., 2011). Normal Spo11-DSB formation requires recruitment of Spo11 and accessory proteins to the meiotic axis (Panizza et al., 2011; Prieler et al., 2005), and Red1/Hop1 are also central to mechanisms that direct Spo11-DSB repair towards use of the homolog as a recombination partner (Carballo etal., 2008; Niu et al., 2005; Schwacha and Kleckner, 1997). Other eukaryotes contain Hop1 analogs that share a domain, called the HORMA domain (Rosenberg and Corbett, 2015), and correlations between these meiotic axis proteins and DSB formation are observed in fission yeast, nematodes and in mammals (Fowler et al., 2013; Goodyer et al., 2008; Wojtasz et al., 2009). Thus, most meiotic interhomolog recombination occurs in the context of a specialized chromosome structure and requires components of that structure.

Meiotic recombination pathways diverge after DSB formation and homolog-directed strand invasion. In budding yeast, about half of events form NCOs via synthesis-dependent strand annealing, a mechanism that does not involve stable recombination intermediates (Allers and Lichten, 2001a; Martini et al., 2011; McMahill et al., 2007) and is suggested to be the predominant HR pathway in mitotic cells (Bzymek et al., 2010; McGill et al., 1989; Mitchel et al., 2013). Most of the remaining events are repaired by a meiosis-specific CO pathway, in which an ensemble of meiotic proteins, called the ZMM proteins, stabilize early recombination intermediates and promote their maturation into double Holliday junction joint molecules (JMs) (Allers and Lichten, 2001a; Börner et al., 2004; Lynn et al., 2007; Schwacha and Kleckner, 1994). These ZMM-stabilized JMs are subsequently resolved as COs (Sourirajan and Lichten, 2008) through the action of the MutLγ complex, which contains the Mlh1, Mlh3, and Exo1 proteins (Argueso et al., 2002; 2004; Khazanehdari and Borts, 2000; Wang et al., 1999; Zakharyevich et al., 2010; 2012). MutLγ does not appear to make significant contributions to mitotic COs (Ira et al., 2003; Welz-Voegele et al., 2002). A minority of events form ZMM-independent JMs that are resolved as both COs and NCOs by the structure-selective nucleases (SSNs) Mus81-Mms4, Yen1, and Slx1-Slx4, which are responsible for most JM resolution during mitosis (Argueso et al., 2004; de los Santos etal., 2003; De Muyt et al., 2012; Zakharyevich et al., 2012) (Ho et al., 2010; Muñoz-Galván et al., 2012) [reviewed by (Wyatt and West, 2014)]. A similar picture, with MutLγ forming most meiotic COs and SSNs playing a minor role, is observed in several other eukaryotes (Falque et al., 2009; Franklin et al., 2006; Hassold et al., 2009; Kochakpour and Moens, 2008; Lhuissier etal., 2007; Plug et al., 1998; Tease and Hultén, 2004) (Higgins et al., 2008; Holloway etal.,2008).

To better understand the factors that promote the unique biochemistry of CO formation during meiosis, in particular MutLγ-dependent JM resolution, we considered two different hypotheses. In the first, expression of meiosis-specific proteins and the presence of high levels of Spo11-DSBs results in nucleus-wide changes in recombination biochemistry, shifting the balance towards MutLγ-dependent resolution of JMs, wherever they might occur. In the second, local features of meiotic chromosome structure, in particular enrichment for meiosis-specific chromosome axis proteins, provides an *in cis* structural environment that favors MutLγ-dependent JM resolution. However, because Spo11-DSBs form preferentially in Red1/Hop1-enriched regions, and because these proteins are required for efficient Spo11-DSB formation and interhomolog repair, it is difficult to distinguish these two models by examining Spo11-initiated recombination alone.

To test these two hypotheses, we developed a system in which meiotic recombination is initiated by the sequence- and meiosis-specific *VMA1* derived endonuclease, VDE (Gimble and Thorner, 1992; Nagai et al., 2003). VDE initiates meiotic recombination at similar levels wherever its recognition sequence *(VRS)* is inserted (Fukuda et al., 2008; Neale et al., 2002; Nogami et al., 2002). VDE-catalyzed DSBs (hereafter called VDE-DSBs) form independent of Spo11 and meiotic axis proteins. However, like Spo11-DSBs, VDE-DSBs form after pre-meiotic DNA replication and are repaired using the same end-processing and strand invasion activities that repair Spo11-DSBs (Fukuda et al., 2003; Hodgson et al., 2011; Neale et al., 2002). We examined resolvase contributions to VDE-initiated CO formation, and obtained evidence that local enrichment for meiotic axis proteins promotes MutLγ-dependent CO formation; while recombination that occurs outside of this specialized environment forms COs by MutLγ-independent mechanisms. We also show that CO formation at a locus, and in particular MutLγ-dependent CO formation, requires Spo11-DSB formation elsewhere in the genome.

## Results

### Using VDE to study meiotic recombination at “hot” and “cold” loci

The recombination reporter used for this study contains a VDE recognition sequence *(VRS)* inserted into a copy of *the ARG4* gene on one chromosome, and an uncleavable mutant recognition sequence *(VRS103)* on the homolog (Figure 1). Restriction site polymorphisms at flanking *HindI*II sites, combined with the polymorphic *VRS* site, allow differentiation of parental and recombinant DNA molecules. This recombination reporter was inserted at two loci: *HIS4* and *URA3,* which are “hot” and “cold”, respectively, for Spo11-initiated recombination and Red1/Hop1 occupancy (Borde et al., 1999; Buhler et al., 2007; Pan et al., 2011; Panizza et al., 2011; Wu and Lichten, 1995); also see Figure 4A and Figure 4—figure supplement 1, below). Consistent with previous reports, Spo11-DSBs and the resulting crossovers, are five times more frequent in inserts at *HIS4* than at *URA3* (Figure 1—figure supplement 1A). When VDE is expressed, ~90% of *VRS* sites at both loci were cleaved by 7 h after initiation of sporulation (Figure 2A), consistent with previous reports that VDE cuts very efficiently (Johnson et al., 2007; Neale et al., 2002; Terentyev et al., 2010). DSBs appeared and disappeared with similar timing at the two loci (Figure 2B), with measures of insert recovery (Figure 2—figure supplement 1A) and levels of interhomolog recombinants relative to cumulative VDE-DSB levels (Figure 2—figure supplement 1B) indicating that ~70% of VDE DSBs are repaired by interhomolog recombination. The remaining *VRS-*containing inserts appear to be lost, consistent with high levels of VDE activity preventing recovery of inter-sister recombinants. Thus, the two VDE recombination reporter inserts undergo comparably high levels of meiotic recombination initiation, regardless of the local intrinsic level of Spo11-initiated recombination.

**Figure 1:**
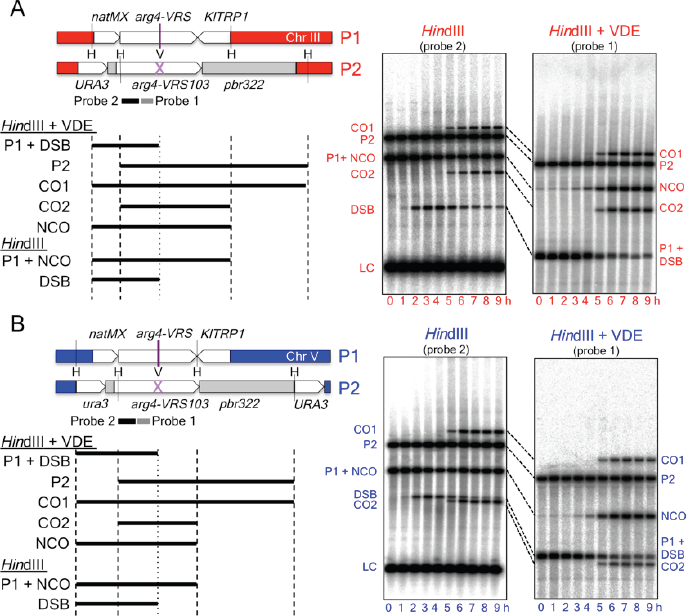
Inserts used to monitor VDE-initiated meiotic recombination. The *HIS4* and *URA3* loci are denoted throughout this paper in red and blue, respectively, and are in Red1/Hop1 enriched and depleted regions, respectively (see Figure 4A and Figure 4— figure supplement 1, below). (**A**) Left—map of VDE-reporter inserts at *HIS4,* showing digests used to detect recombination intermediates and products; right—representative Southern blots. One parent contains *ARG4* sequences with a VDE-recognition site *(arg4-VRS),* flanked by an noursethricin-resistance module *(natMX,* Goldstein and McCusker, 1999) and the *Kluyveromyces lactis TRP1* gene *(KlTRP1,* Stark and Milner, 1989); the other parent contains *ARG4* sequences with a mutant, uncuttable *VRS* site *(arg4-VRS103,* Nogami et al., 2002) flanked by *URA3* and pBR322 sequences. Digestion with *HindIII* (H) and VDE (V) allows detection of crossovers (CO1 and CO2) and noncrossovers (NCO); digestion with *HindI*II alone allows detection of crossovers and DSBs. The *Hin*dIII-alone blot has been probed with a fragment (probe 2) that hybridizes to the insert loci and to the native *ARG4* locus on chromosome VIII; this latter signal serves as a loading control (LC). Times after induction of meiosis that each sample was taken are indicated below each lane. (**B**) map of VDE-reporter inserts at *URA3* and representative Southern blots; details as in (A). Strain, insert and probe details are given in Materials and Methods and Supplementary Table 1.

**Figure 2.**
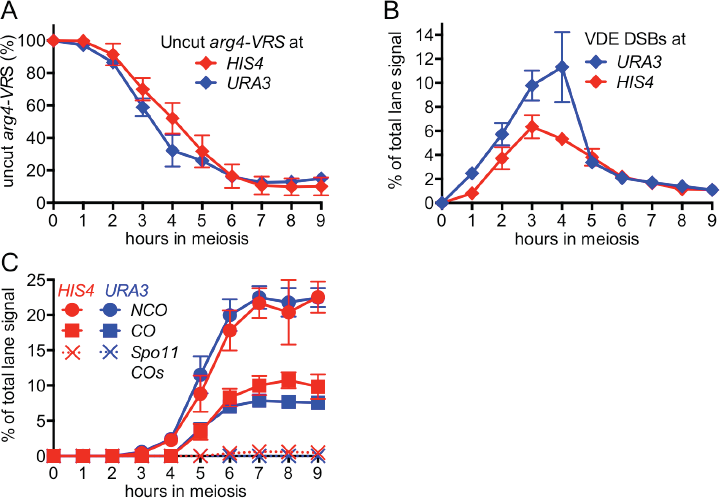
VDE-initiated recombination occurs at similar levels at the two insert loci. (**A**) Cumulative DSB levels are similar at the two insert loci. The fraction of uncut VRS-containing chromosomes (Parent 1) was determined by subtracting the amount of the NCO band in *HindIU* + VDE digests from the amount of the Parent 1 + NCO band in *HindIU* digests. (**B**) Non-cumulative VDE-DSB frequencies, measured as fraction of total lane signal in *HindIU* digests, excluding loading controls. (**C**) Crossover (average of CO1 and CO2) and noncrossover frequencies, measured in *HindIII-VDE* digests. Solid lines—recombinants from cells expressing VDE; dashed lines—Spo11-initiated crossovers from *vde* strains, see also Figure 1—figure supplement 1. Values are the average of two independent experiments; error bars represent standard error of mean. For COs, error is calculated for the average values of CO1 and CO2 from both experiments. Representative Southern blots are shown in Figure 1 and Figure 1—figure supplement 1C.

When VDE-DSBs are repaired by interhomolog recombination, *VRS* sequences are converted to *VRS103,* and become resistant to digestion by VDE. We therefore used *Hin*dIII/VDE double digest to score VDE-initiated recombination (Figure 1). By comparing levels of recombinants in VDE-expressing and *vdeΔ* strains, we determined that Spo11-initiated events make a negligible contribution to recombinants scored in VDE-expressing strains (Figure 2C, Figure 1—figure supplement 1). VDE-initiated recombinants formed at high frequencies at both *HIS4* and *URA3* (Figure 2C), and NCOs exceeded COs by approximately twofold at *HIS4* and threefold at *URA3* (Figure 2D). These values are within the range observed in genetic studies of Spo11-induced gene conversion in budding yeast (Fogel et al., 1979), but differ from the average of near-parity between NCOs and COs observed in molecular assays (Lao et al., 2013; Martini et al., 2006). VDE, unlike Spo11, frequently cuts both sister chromatids (Gimble and Thorner, 1992; Zhang et al., 2011), and this may reduce the fraction of DSBs that are repaired as COs (Malkova et al., 2000).

### MutLγ makes different contributions to VDE-initiated CO formation at the two insert loci

While VDE-initiated recombination occurred at similar levels in inserts located at *HIS4* and at *URA3,* we observed a marked difference between the two loci, in terms of the resolvase-dependence of CO formation (Figure 3). At the *HIS4 locus,* COs were reduced in *mlh3* mutants, which lack MutLγ, by ~60% relative to wild type. COs were reduced by ~30% in in *mms4-mdyen1Δ slx1Δ* mutants, (hereafter abbreviated as *ssn* mutants), which lack the three structure selective nucleases (SSNs) active during both meiosis and the mitotic cell cycle, and by ~75% in *mlh3 ssn* mutants. Thus, like Spo11-initiated COs, VDE-initiated COs in inserts at *HIS4* are primarily MutLγ-dependent, and less dependent on SSNs. In contrast, COs in inserts located at *URA3* were reduced by only ~10% in *mlh3,* by ~40% in *ssn* mutants, and by ~60% in *mlh3 ssn* mutants, leaving approximately the same residual CO levels as was seen at *HIS4.* Thus, SSNs make a substantially greater contribution to VDE-initiated CO formation at *URA3* than does MutLγ, and MutLγ’s contribution becomes substantial only in the absence of SSNs.

**Figure 3.**
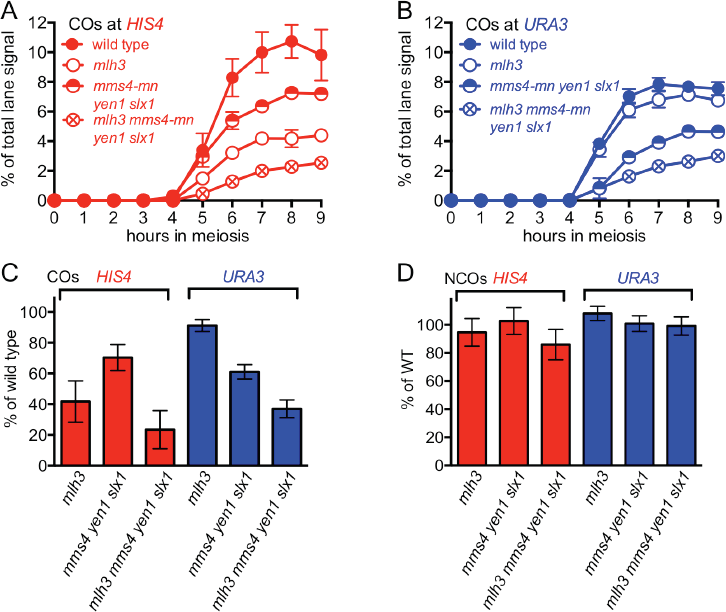
Different resolvase-dependence of crossover formation at the two insert loci. (**A**) Crossover frequencies (average of CO1 and CO2) measured as in Figure 2(C) from *HIS4* insert-containing mutants lacking MutLγ *(mlh3),* structure-selective nucleases *(mms4-mn yen1 slx1)* or both resolvase activities *(mlh3 mms4-mn yen1 slx1).* (**B**) Crossover frequencies in *URA3* insert-containing strains, measured as in panel A. Values are the average of two independent experiments; error bars represent standard error of mean (SEM). (**C**) Final crossover levels (average of 8 and 9 h values), expressed as percent of wild type. Note that, in *mlh3* mutants, crossovers in *HIS4* inserts are reduced more than twofold, while crossovers in *URA3* inserts are reduced by less than 10%. (**D**) Final noncrossover levels (average of 8 and 9 h values), expressed as percent of wild type. Percent of wild type (WT) values are calculated from average values of two independent experiments; error bars are calculated as (SEM_wr_/Average_wr_) + (SEM_mutant_/Average_mutant_), normalized to 100%. Representative Southern blots are in Figure 3—figure supplement 2.

At both insert loci, *ssn* and *mlh3 ssn* mutants accumulated DNA species with reduced electrophoretic mobility (Figure 3—figure supplement 2). These slower-migrating species contain branched DNA molecules, as would be expected for unresolved joint molecules (D. M., unpublished observations). Steady state VDE-DSB levels and final NCO levels were similar in all strains (Figure 3D, Figure 3—figure supplement 1), indicating that resolvases do not act during the initial steps of DSB repair, and that most meiotic NCOs form by mechanisms that do not involve Holliday junction resolution (Allers and Lichten, Muyt et al., 2012; Sourirajan and Lichten, 2008; Zakharyevich et al., 2012).

### Altered Hop1 occupancy in *pch2* mutants is associated with altered MutLγ-dependence of VDE-initiated COs

The marked MutLγ-dependence and -independence of VDE-initiated COs in inserts at *HIS4* and at *URA3,* respectively, are paralleled by levels of occupancy at the two loci of the meiotic axis proteins, Hop1 and Red1 (Panizza et al., 2011); Figure 4A, Figure 4—figure supplement 1A). To ask if differential Hop1 occupancy is responsible for the observed differences in CO formation at these loci, we examined the resolvase-dependence of VDE-initiated COs in *pch2Δ* mutants. Pch2 is a conserved AAA ATPase that maintains the nonuniform pattern of Hop1 occupancy along meiotic chromosomes (Börner et al., 2008; Joshi et al., 2009). The different Hop1 occupancies seen in wild type were preserved early in meiosis in *pch2Δ* mutants (Figure 4A, Figure 4—figure supplement 1A), consistent with previous findings that, in *pch2* cells, Spo11-DSB patterns are not altered in most regions of the genome (Vader et al., 2011). By contrast, at later times (4-5 h after initiation of meiosis), *pch2Δ* mutants displayed reduced Hop1 occupancy at *HIS4,* more closely approaching the lower occupancy levels seen throughout meiosis at *URA3* (Figure 4A; Figure 4—figure supplement 4A).

**Figure 4.**
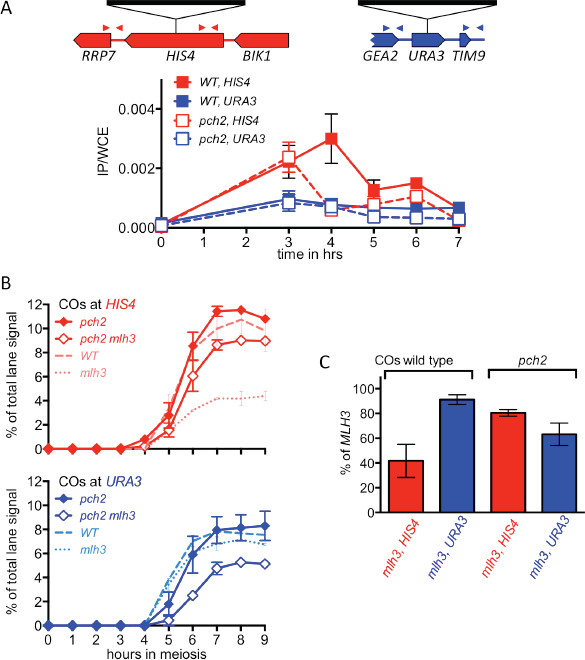

*pch2Δ* mutants display altered Hop1 occupancy and crossover MutLγ-dependence. (**A**) Hop1 occupancy at insert loci, determined by chromatin immunoprecipitation and quantitative PCR. Top—cartoon of insert loci, showing the location of primer pairs used. Bottom—relative Hop1 occupancy, expressed as the average ratio of immunopreciptiate/input extract for both primer pairs (see Materials and Methods for details). Values are the average of two independent experiments; error bars represent standard error of the mean. (**B**) VDE-initiated CO frequencies measured as in Figure 2(C) at *HIS4* (top) and *URA3* (bottom) in *pch2Δ* (solid diamonds) and *pch2Δ mlh3Δ* (open diamonds). Crossovers from wild type (dashed line) and *mlhΔ* (dotted line) from Figure 3 are shown for comparison. Values are from two independent experiments; error bars represent standard error of the mean. Representative Southern blots are in Figure 4— figure supplement 2. (**C**) Extent of CO reduction in *mlhΔ* mutants, relative to corresponding *MLH3* strains. *PCH2* genotype is indicated at the top; values and error bars are calculated as in Figure 3(C).

While VDE-induced DSB dynamics and NCO levels were similar in *PCH2* and *pch2Δ* strains (Figure 4—figure supplement 1B, C), the loss of Pch2 was accompanied by a marked change in MutLγ contributions to VDE-initiated COs. The majority of COs became MutLγ-independent at both insert loci (Figure 4B, C). At *HIS4,* the fraction of COs that were MutLγ-dependent decreased substantially (from ~60% in *PCH2* to 20% in *pch2Δ),* while at *URA3* the fraction that were MutLγ-dependent increased (from ~10% to 37%). Thus, in *pch2Δ* mutants, the similarity of Hop1 occupancy at later times in meiosis is paralleled by a shift in the MutLγ-dependence of VDE-initiated COs, with contributions of MutLγ to COs in inserts at *HIS4* and *URA3* becoming more similar.

### Spo11-DSBs promote VDE-initiated, MutLγ-dependent COs

All experiments reported above used cells with wild-type levels of Spo11-DSBs. While VDE-DSBs form at similar levels and timing in *SPO11* and *spo11* mutant cells (Johnson et al., 2007; Neale et al., 2002; Terentyev et al., 2010), features of VDE-DSB repair, including the extent of end resection, are strongly influenced by the presence or absence of Spo11-DSBs (Neale et al., 2002). To determine if other aspects of VDE-initiated recombination are also affected, we examined VDE-initiated recombination in a catalysis-null *spo11-Y135F*mutant, hereafter called *spo11.* In *spo11* mutants, VDE-DSB dynamics and NCO formation were similar in inserts at *HIS4* and *URA3,* were comparable to those seen in wild type (Figure 5— figure supplement 1), and were independent of HJ resolvase activities (Figure 5—figure supplement 1). In contrast, the absence of Spo11-DSBs substantially reduced VDE-induced COs, resulting in virtually identical CO timing and levels at the two loci (Figure 5A). Unlike the ~60% reduction in COs seen at *HIS4* in *SPO11 mlhΔ* (Figure 3C), final CO levels were similar in *spo11 mlhΔ* and *spo11 MLH3* strains, at both *HIS4* and *URA3,* and similar CO reductions were observed at both loci in *spo11 snn* mutants (Figure 5B, C). Thus, processes that depend on Spo11-DSBs elsewhere in the genome are important to promote VDE-initiated COs, and appear to be essential for MutLγ-dependent CO formation.

**Figure 5.**
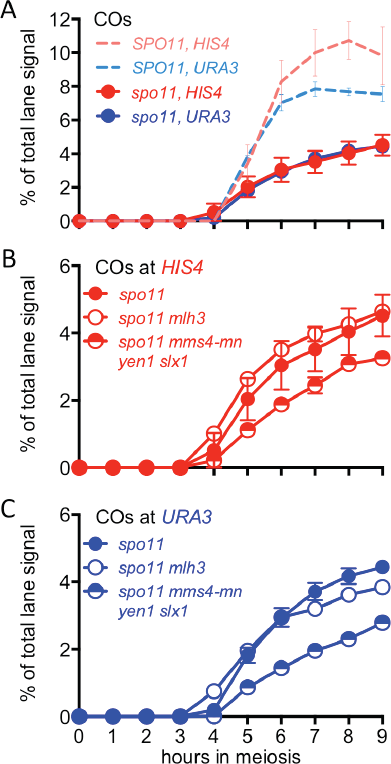

VDE-initiated COs are reduced and are Mutiny-independent in the absence of Spoil activity. (**A**) VDE-initiated crossover frequencies as measured in Figure 2(C) in *spoll-Y135F*strains (dark solid lines) in inserts at *HIS4* (red) and at *URA3* (blue). Data from the corresponding *SPO11* strains (dotted lines, from Figure 2C) are presented for comparison. (**B**) CO frequencies in *HIS4* inserts in *spo11* strains that are otherwise wild-type *(spo11)* or lack either MutLγ *(spo11 mlhΔ)* or structure-selective nucleases *(spo11 mms4-mnyenlA slxlA).* (**C**) As in B, but with strains with inserts at *URA3.* Values are from two independent experiments; error bars represent standard error of the mean.

## Discussion

### Local chromosome structure influences meiotic CO formation

We examined the contribution of different Holliday junction resolvases to VDE-initiated CO-formation in recombination reporter inserts at two loci, *HIS4* and *URA3,* which are “hot” and “cold”, respectively, for Spo11-inititiated recombination and for occupancy by the meiotic chromosome axis proteins, Hop1 and Red1. VDE-initiated COs at *HIS4* are similar to those initiated by Spo11, in that most depend on MutLγ. In contrast, VDE-initiated COs at the “cold” locus, *URA3,* more closely resemble mitotic COs, which are independent of MutLγ, but are substantially dependent on SSNs (Ho et al., 2010; Ira et al., 2003; Muñoz-Galván et al., 2012; Welz-Voegele et al., 2002). Locus-dependent differences in MutLγ-dependence are reduced in *pch2Δ* mutants, as are differences in Hop1 occupancy at later times in meiosis I prophase. Based on these findings, we suggest that local chromosome context exerts an important influence on the biochemistry of CO formation during meiosis, and that factors responsible for creating DSB-hot and -cold domains also create corresponding domains where different DSB repair pathways are dominant. An attractive hypothesis (Figure 6) is that regions enriched for meiosis-specific axial element proteins create a chromosomal environment that promotes meiotic DSB formation, limits inter-sister recombination, preferentially loads ZMM proteins (Joshi et al., 2009; Serrentino et al., 2013), and is required for recruitment of MutLγ. In such regions, where most Spo11-dependent events occur, recombination intermediates will have a greater likelihood of being captured by axis-associated ZMM proteins, and consequently being resolved as COs by MutLγ. Regions with lower axial element protein enrichment are less likely to recruit ZMM proteins and MutLγ; DSB repair and CO formation in these regions is more likely to involve non-meiotic mechanisms. In short, the meiotic genome can be thought of as being apportioned between two different environments: meiotic axis protein-enriched regions, where “meiotic” recombination pathways predominate; and meiotic axis protein-depleted regions, in which recombination events more closely resemble those seen in mitotic cells.

**Figure 6.**
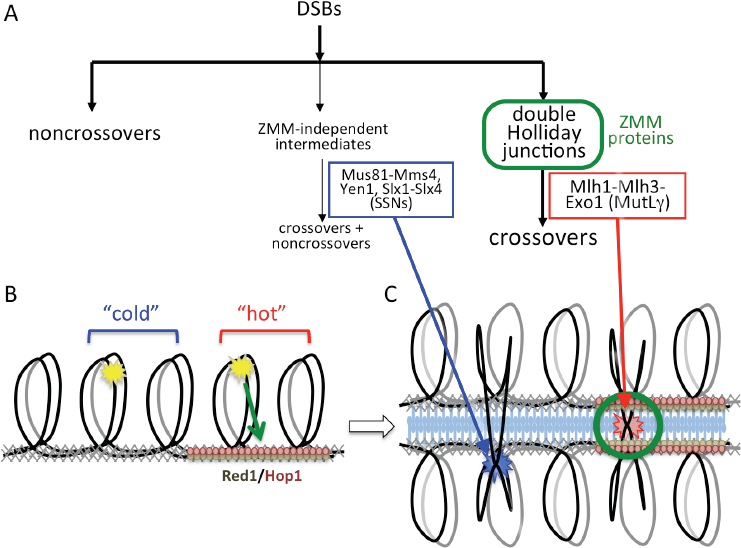

Different resolvase functions in different genome domains. (**A**) Early crossover decision model for meiotic recombination (Bishop and Zickler, 2004; Hollingsworth and Brill, 2004) illustrating early noncrossover formation, a major pathway where recombination intermediates form in the context of ZMM proteins and are resolved by MutLγ to form crossovers, and a minor pathway where ZMM-independent intermediates are resolved by SSNs as both crossovers and noncrossovers. (**B**) Division of the meiotic genome into meiotic axis-protein-enriched “hot” domains (red) that are enriched for Red1 and Hop1, and “cold” domains where Red1 and Hop1 are depleted. VDE DSBs (yellow stars) can be directed to form efficiently in either domain, but only VDE DSBs that form in “hot” domains can be recruited to the meiotic axis. (**C**) DSBs in “hot” domains can form joint molecules (red star) in the context of ZMM proteins and the synaptonemal complex, and thus can be resolved by MutLγ-dependent activities. DSBs in “cold” domains form joint molecules outside of this structural context, and are resolved by MutLγ-independent activities.

While the current study is the first to directly query the effect of chromosome context on JM resolution, others have obtained results that are consistent with an effect of local chromosome context on meiotic DSB repair. Malkova and coworkers used the HO endonuclease to initiate recombination in meiotic cells at *LEU2,* also a “hot” locus (Panizza et al., 2011; Wu and Lichten, 1995). The resulting COs were dependent on Msh4, a ZMM protein, to the same degree as are Spo11-induced COs, suggesting that these nuclease-induced COs at the axis enriched *LEU2* locus (Panizza et al., 2011) were the products of ZMM/MutLγ-dependent JM resolution (Malkova et al., 2000). Consistent with our suggestion that different recombination mechanisms operate in different parts of the genome, the meiotic genome also appears to be divided into regions that respond to DNA damage in different ways. Treatment of meiotic yeast cells with phleomycin, a DSB-forming agent, triggers Rad53 phosphorylation, as it does in mitotic cells, while Spo11-DSBs do not (Cartagena-Lirola et al., 2008). This indicates that Spo11-DSBs form in an environment that is refractory to Rad53 recruitment and modification, but that there also are regions in the meiotic genome where exogenously-induced damage can trigger the mitotic DNA damage response. In light of these ideas, it is interesting that the meiotic defects of *spo11* mutants in a variety of organisms are often only partially rescued by treatment with exogenous agents that cause DSBs (Bowring et al., 2006; Celerin et al., 2000; Dernburg et al., 1998; Loidl and Mochizuki, 2009; Pauklin et al., 2009; Storlazzi et al., 2003; Thorne and Byers, 1993). While other factors may be responsible for the limited rescue observed, we suggest that it reflects the random location of exogenously-induced DSBs, with only a subset forming in regions where repair is likely to form interhomolog COs that promote proper homolog segregation.

### The interplay of resolvase activities is chromosome context-dependent

Although we observe marked differences in the contributions of different resolvases to VDE-induced CO formation at *HIS4* and at *URA3,* there is no absolute demarcation between MutLγ and SSN activities at the two loci. At *HIS4,* where MutLγ predominates, *ssn* mutants still display a modest reduction in VDE-initiated COs when MutLγ is active, but an even greater relative reduction in the absence of MutLγ. These findings are consistent with previous studies suggesting that, in the absence of MutLγ, SSNs are able to serve as a backup JM resolvase (Argueso et al., 2004; De Muyt et al., 2012; Zakharyevich et al., 2012). Our current data indicate that the converse may also be true, since at *URA3,* MutLγ appears to make a greater contribution to CO formation in the absence of SSNs than in their presence. However, in our studies, JMs are more efficiently resolved in *mlhΔ* mutants than in *ssn* mutants, which display persistent unresolved JMs. Therefore, if MutLγ acts as a back-up resolvase, it can do so in only a limited capacity, possibly reflecting a need for a specific chromosome structural context in which to function efficiently. The absence of such a meiosis-specific chromosome context may explain why MutLγ does not appear to contribute to CO formation during the mitotic cell cycle (Ira et al., 2003; Welz-Voegele et al., 2002), although the lower expression of *MLH3* expression in mitotic cells (Brar et al., 2012; Primig et al., 2000) may also contribute.

Both VDE-induced and Spo11-induced COs form at significant frequencies in *mlhΔ ssn* mutants, which lack all four of the HJ resolvase activities thought to function during meiosis (Figure 3, see also (Argueso et al., 2004; Zakharyevich et al., 2012). These residual crossovers may reflect the activity of a yet-unidentified JM resolvase; they may also reflect the production of half-crossovers by break-induced replication (Ho et al., 2010; Kogoma, 1996; Llorente et al., 2008) or by other mechanisms that do not involve dHJ-JM formation and resolution (Ho et al., 2010; Ivanov and Haber, 1995; Muñoz-Galván et al., 2012; Prado and Aguilera, 1995).

### Genome-wide Spo11-DSBs promote VDE-initiated COs and are required for chromosome context-dependent differentiation of VDE DSB repair

In catalysis-null *spo11-Y135F*mutants, most VDE-DSBs are repaired by interhomolog recombination (Figure 4, Figure 4—figure supplement 2), indicating that a single DSB can efficiently search the meiotic nucleus for homology. However, VDE-promoted COs are substantially reduced in *spo11* mutants (Figure 4), as has been observed with HO endonuclease-induced meiotic recombination (Malkova et al., 2000). Moreover, in *spo11* mutants, virtually all VDE-initiated COs are MutLγ-independent (Figure 4, Figure 4—figure supplement 2). Because Hop1s occupancy of cohesin sites is not noticeably altered in *spo11-Y135F*mutants (Franz Klein, personal communication), these findings indicate that, in addition to the local effects of meiotic chromosome structure suggested above, CO formation is affected by processes that require Spo11-DSBs elsewhere in the genome.

Meiotic DSB repair occurs concurrently with homolog pairing and synapsis (Börner et al., 2004; Padmore et al., 1991), and efficient homolog synapsis requires wild-type DSB levels, indicating that multiple interhomolog interactions along a chromosome are needed for stable homolog pairing (Henderson and Keeney, 2004). To account for the reduced levels and MutLγ-independence of VDE-initiated COs in *spo11* mutants, we suggest that a single VDE-DSB is not sufficient to promote stable homolog pairing, and that additional DSBs along a chromosome are needed to promote stable homolog pairing, which in turn is needed to form ZMM protein-containing structures that stabilize JMs and recruit MutLγ. However, the 140-190 Spo11-DSBs that form in each meiotic cell (Buhler et al., 2007; Martini et al., 2011; Pan et al., 2011) are also expected to induce a nucleus-wide DNA damage-response, and to compete with other DSBs for repair activities whose availability is limited, and both have the potential to alter recombination biochemistry at VDE-DSBs (Johnson et al., 2007; Neale et al., 2002). Thus, while we believe it likely that defects in homolog pairing and synapsis are responsible for the observed impact of *spo11* mutation on VDE-initiated CO formation, it remains possible that it is due to changes in DNA damage signaling, repair protein availability, or in other processes that are affected by global Spo11-DSB levels.

### Concluding remarks

We have provided evidence that structural features of the chromosome axis, in particular the enrichment for meiosis-specific axis proteins, create a local environment that directs recombination to “meiotic” biochemical pathways. In the remainder of the genome, biochemical processes more typical of mitotic recombination function. In other words, the transition to meiosis from the mitotic cell cycle does not require a global inhibition of mitotic” recombination mechanisms, which remain active in the meiotic nucleus and have the capacity to act in recombination events that occur outside of the local “meiotic “ structural context. It is well established that this local chromosome context influences the first step in meiotic recombination, Spo11-catalyzed DSB formation (Panizza et al., 2011; Prieler et al., 2005). Our work shows that it also influences the last, namely the resolution of recombination intermediates to form COs. It will be of considerable interest to determine if other critical steps in meiotic recombination, such as choice between sister and homolog as a DSB repair partner, are also influenced by local aspects of chromosome structure.

In the current work, we focused on correlations between local enrichment for the meiosis-specific axis protein Hop1 and Holliday junction resolution activity during CO formation. Other HORMA domain proteins, including HIM-3 and HTP-1/2/3 in *C. elegans,* ASY3 in *A thaliana* and HORMAD1/2 in *M. musculus* also have been reported to regulate recombination and homolog pairing (Kim et al., 2014; Martinez-Perez and Villeneuve, 2005) (Ferdous et al., 2012; Fukuda et al., 2010; Wojtasz et al., 2009), suggesting that HORMA domain proteins may provide a common basis for the chromosome context-dependent regulation of meiotic recombination pathways in eukaryotes.

## Materials and Methods

### Yeast strains

All yeast strains are of SK1 background (Kane and Roth, 1974), and were constructed by standard genetic crosses or by direct transformation. Genotypes and allele details are given in Supplementary Table 1. Recombination reporter inserts with *arg4-VRS103* contain a 73nt *VRS103* oligonucleotide containing the mutant VDE recognition sequence from the *VMA1-103* allele (Fukuda et al., 2007; Nogami et al., 2002) inserted at the *EcoRV* site in *ARG4* coding sequences within a pBR322-based plasmid with *URA* and *ARG4* sequences, inserted at the *URA3* and *HIS4* loci, as described (Wu and Lichten, 1995). Recombination reporter inserts with the cleavable *arg4-VRS* (Neale et al., 2002) were derived from similar inserts but with flanking repeat sequences removed, to prevent repair by single strand annealing (Pâques and Haber, 1999). This was done by replacing sequences upstream and downstream of *ARG4* with *natMX* (Goldstein and McCusker, 1999) and *K. lactis TRP1* sequences (Stark and Milner, 1989) respectively (see Supplementary Table 1 legend for details. The resulting *arg4-VRS* and *arg4-VRS103* inserts share 3.077 kb of homology.

VDE normally exists as an intein in the constitutively-expressed *VMA1* gene (Gimble and Thorner, 1993), resulting in low levels of DSB formation in presporulation cultures (data not shown), probably due to small amounts VDE incidentally imported to the nucleus during mitotic growth (Nagai et al., 2003). To further restrict VDE DSB formation, strains were constructed in which *VDE* expression was copper-inducible. These strains contain the *VMA1-103* allele (Nogami et al., 2002), which provides wild type *VMA1* function, but lacks the VDE intein and is resistant to cleavage by VDE. To make strains in which *VDE* expression was copper-inducible, *VDE* coding sequences on an *Eco*RI fragment from pY2181 (Nogami et al., 2002) (a generous gift from Drs. Satoru Nogami and Dr. Yoshikazu Ohya) were inserted downstream of the CUP1 promoter in plasmid pHG40, which contains the *kanMX* selectable marker and a ~1kb *CUP1* promoter fragment (Jin et al., 2009), to make pMJ920, which was then integrated at the *CUP1* locus.

### Sporulation

Yeast strains were grown in buffered liquid presporulation medium and shifted to sporulation medium as described (Goyon and Lichten, 1993), except that sporulation medium contained 10uM CuSO4 to induce *VDE* expression. All experiments were performed at 30°C. Independent experiments were performed either on different days, or on the same day with cultures derived from independent single colonies.

### DNA extraction and analysis

Genomic DNA was prepared as described (Allers and Lichten, 2000). Recombination products were detected on Southern blots containing genomic DNA digested with *HindI*II and *VDE (PI-SceI,* New England Biolabs), using specific buffer for *PI-SceI.* Samples were heated to 65° C for 15 min before loading to disrupt VDE-DNA complexes; gels contained 0.5% agarose in 45 mM Tris Borate + 1 mM EDTA (1X TBE) and were run at 2 V/cm for 24-25 hours. DSBs were similarly detected on Southern blots, but were digested with *HindI*II alone as previously described (Goldfarb and Lichten, 2010), and electrophoresis buffer was supplemented with 4mM MgCl2. Gels were transferred to membranes and hybridized with radioactive probe as described (Allers and Lichten, 2001a; 2001b), and were imaged and quantified using a Fuji FLA-5100 phosphorimager and ImageGauge 4.22 software. *HindI*II*-VDE* gel blots were probed with *ARG4* sequences from −430 to +63 nt relative to *ARG4* coding sequences (Probe 1, Figure 1). *HindI*II gel blots were probed with sequences from the *DED81* gene (+978 to +1650 nt relative to *DED81* coding sequence), which is immediately upstream of *ARG4* (Probe 2, Figure 2).

### Chromatin immunoprecipitation and quantitative PCR

Cells were formaldehyde-fixed by adding 840 μl of a 36.5-38% formaldehyde solution (Sigma) to 30 ml of meiotic cultures, incubating for 15 minutes at room temperature, and quenched by the addition of glycine to 125 mM. Cells were harvested by centrifugation, resuspended in 500 μl lysis buffer (from (Strahl-Bolsinger et al., 1997) except with 1mg/ml Bacitracin and cOmplete Roche protease inhibitor cocktail (1 tablet/10ml) as protease inhibitors) and lysed at 4°C via 10 cycles of vortexing on a FastPrep24 (MP Medical) at 4 M/sec for 40 secs, with 5 minute pauses between runs. Lysates were then sonicated to yield an average DNA size of 300 bp and clarified by centrifugation at 21,130 RCF for 20 minutes. 1/50th of the sample was removed as input, and 2μl of anti-Hop1 (a generous gift from Nancy Hollingsworth) was added to the remainder (~490 μl) and incubated with gentle agitation overnight at 4°C. Antibody complexes were purified by addition of 20 μl of 50% slurry of Gammabind G Sepharose beads (GE healthcare), with further incubation for 3 hours at 4°C, followed by pelleting at 845 RCF for 30 seconds. Beads were then washed and processed for DNA extraction as described (Blitzblau 1993; Andreas Hochwagen, personal communication).

qPCR analysis of purified DNA from input and immunoprecipitated samples used primer pairs that amplify two regions (chromosome III coordinates 65350-65547 and 68072-68271, Saccharomyces Genome Database release # R64-2-1) flanking the *HIS4* gene, and two regions (chromosome V coordinates 115119-115317 and 117728-117922) flanking the *URA3* gene (see Figure 1—figure supplement 1). Primers and genomic DNA from input and immunoprecipitated samples were mixed with iQ SYBR green supermix (Biorad) and analyzed using a Biorad iCycler.

## Acknowledgements

We thank Robert Shroff, Anuradha Sourirajan, Satoru Nogami, Yoshikazu Ohya, and Nancy Hollingsworth for strains and reagents, Andreas Hochwagen and Franz Klein for communicating unpublished information, and Dhruba Chattoraj, Julie Cooper, and Alex Kelly for comments on the manuscript.

---

The following figure supplement is available for Figure 1: **Figure supplement 1.** Spo11-initiated events at the two insert loci.

**Figure 1—figure supplement 1.**
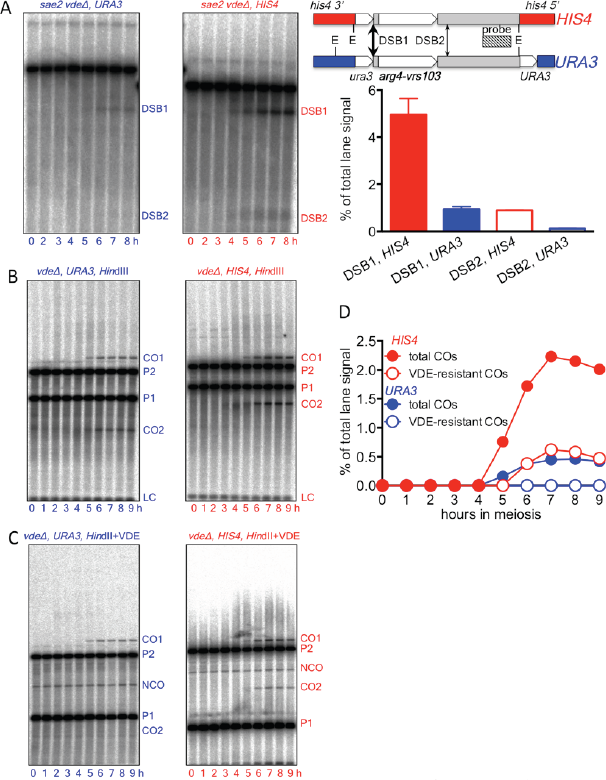

Spo11-initiated events at the two insert loci. (**A**) Spo11-catalyzed DSBs are more frequent in inserts at *HIS4* that at *URA3.* Left—Southern blots of *EcoRI* digests of DNA from strains lacking VDE, probed with pBR322 sequences, showing Spo-DSBs in the Parent 2 insert (see Figure 1, main panel) in resection/repair-deficient *sae2* Δ mutant strains. Right—location of DSBs and probe and DSB frequencies (average of 7 and 8 h samples from a single experiment; error bars represent standard error of mean). (**B**) Southern blots of *Hind*II*I* digests of DNA from strains lacking VDE, to detect total Spo11-initiated crossovers. (**C**) Southern blots of*Hin*dIII-VDE double digests of the same samples, to determine the background contribution of Spo11-initiated COs in subsequent experiments measuring VDE-initiated COs, which will be VDE-resistant due to conversion of the *VRS*site to *VRS103.* (**D**) Quantification of data in panels B (total COs) and C (VDE-resistant COs). Data are from a single experiment.

The following figure supplements is available for Figure 2: **Figure supplement 1.** 70-80% of VDE-DSBs are repaired.

**Figure 2—figure supplement 1.**
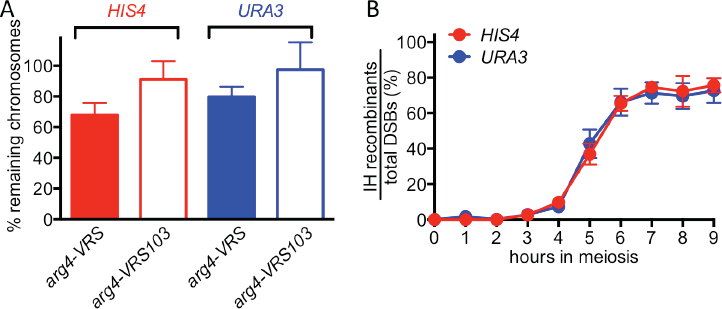

70-80% of VDE-DSBs are repaired. (**A**) Fraction of inserts remaining, calculated using *HindUI* digests (see Figure 1). For the *arg4-VRS103* insert, the ratio (Parent 2 + CO2)/ (0.5 x LC) was calculated at 9 h, and was then normalized to the 0 h value. For the *arg4-VRS* insert, a similar calculation was made: (Parent 1 + NCO + CO1)/(0.5 x LC) (**B**) Relative recovery of interhomolog recombination products, calculated using///*n*dIII-VDE double digests (see Figure 1). The sum of CO (average of CO1 and CO2) and NCO frequencies was divided by the frequency of total DSBs, as calculated in Figure 2A. Data are the average of two independent experiments; error bars represent standard error of mean.

The following figure supplements are available for Figure 3: **Figure supplement 1.**

VDE-DSB and NCO frequencies in resolvase mutants.

**Figure supplement 2.** Representative Southern blots.

**Figure 3—figure supplement 1.**
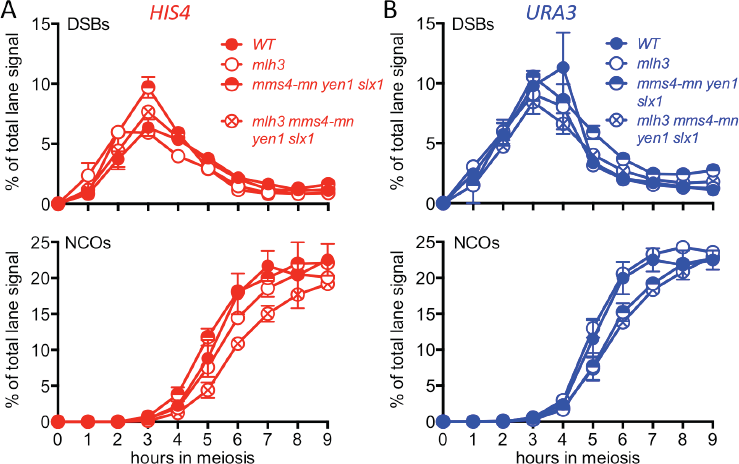

VDE-DSB and NCO frequencies in resolvase mutants. (**A**) VDE-DSB frequencies (top), measured as in Figure 2B, and NCO frequencies (bottom), measured as in Figure 2C, from *HIS4* insert-containing strains. (**B**) As panel A, with strains containing inserts at *URA3.* Data are the average of two independent experiments; error bars represent standard error of mean. Representative Southern blots are in Figure 3— figure supplement 2.

**Figure 3—figure supplement 2.**
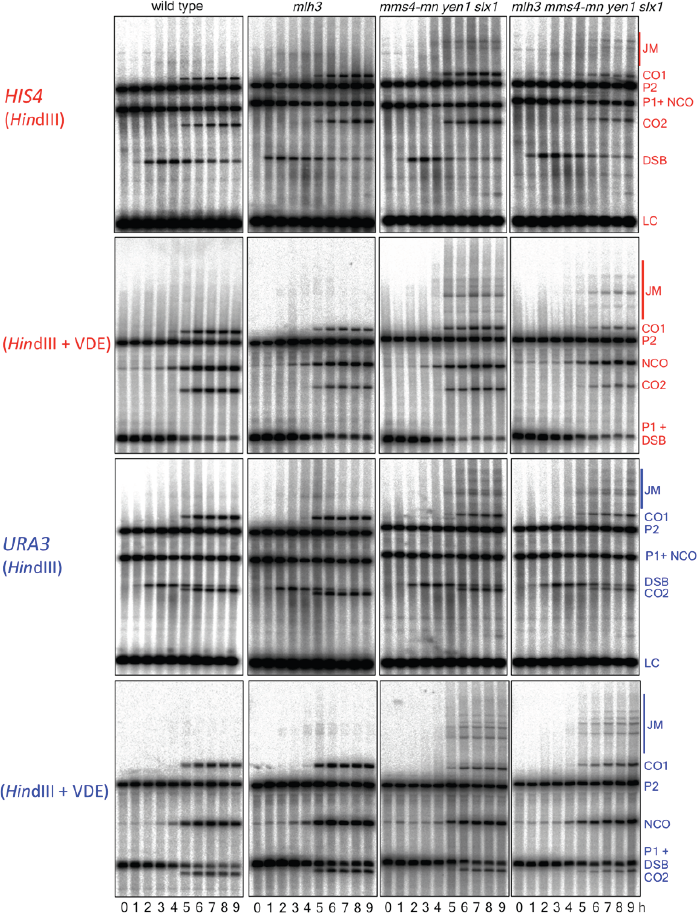

Representative Southern blots. Blots are of *HindIII HindI*II*-VDE* digests of DNA from *HIS4* insert-containing strains (top) and from *URA3* insert-contaning strains (bottom). Gel labels as in Figure 1; JM—joint molecule recombination intermediates.

The following figure supplements are available for Figure 4: **Figure supplement 1.**

Hop1 occupancy, DSBs and NCOs.

**Figure supplement 2.** Representative Southern blots.

**Figure 4—figure supplement 1.**
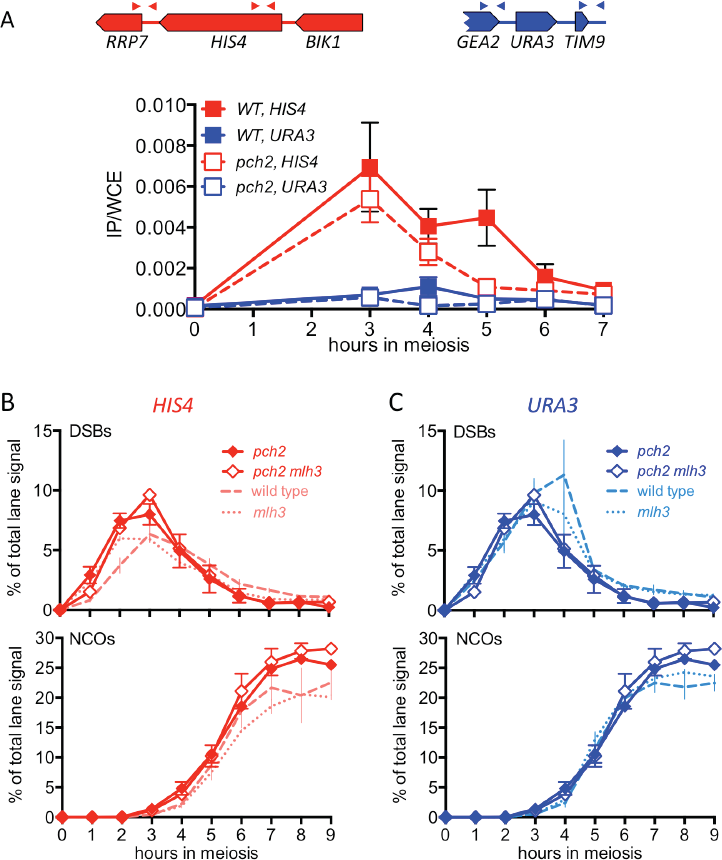

Hop1 occupancy, DSBs and NCOs. (**A**) Hop1 occupancy at corresponding loci lacking inserts, determined as in Figure 4A. Occupancy at *HIS4* is from strains with inserts at *URA3,* and vice versa. (**B**) DSBs and NCO frequencies in inserts at *HIS4,* determined as in Figure 2B and 2C, respectively. Symbols are as in Figure 4B. (**C**) DSBs and NCOs in inserts at *URA3,* details as in panel B. Values are from two independent experiments; error bars represent standard error of the mean. Representative Southern blots are in Figure 4—figure supplement 2.

**Figure 4—figure supplement 2.**
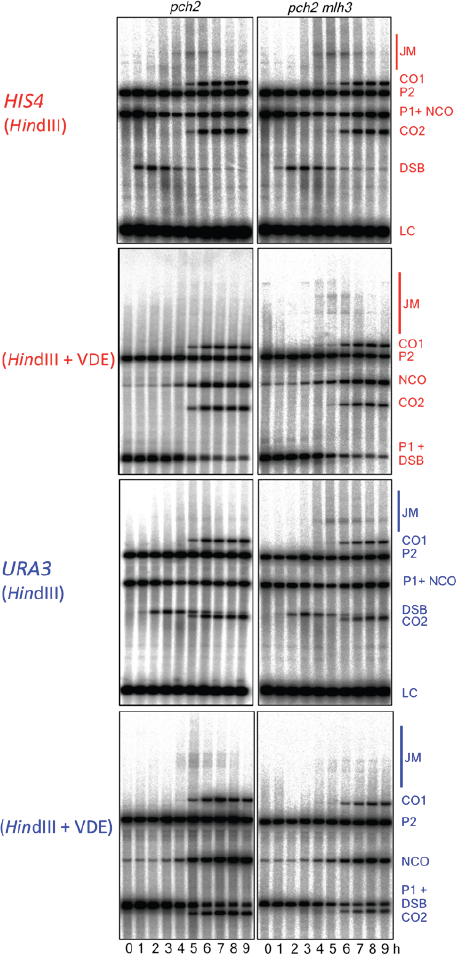

Representative Southern blots. Blots are of *HindIII*; *Hin*dIII-VDE digests of DNA from *HIS4* insert-containing strains (top) and from *URA3* insert-contaning strains (bottom). Gel labels as in Figure 1; JM—joint molecule recombination intermediates.

The following figure supplements are available for Figure 5: **Figure supplement 1**.

DSBs and recombinant products in *spo11* strains.

**Figure supplement 2.** Representative Southern blots.

**Figure 5—figure supplement 1.**
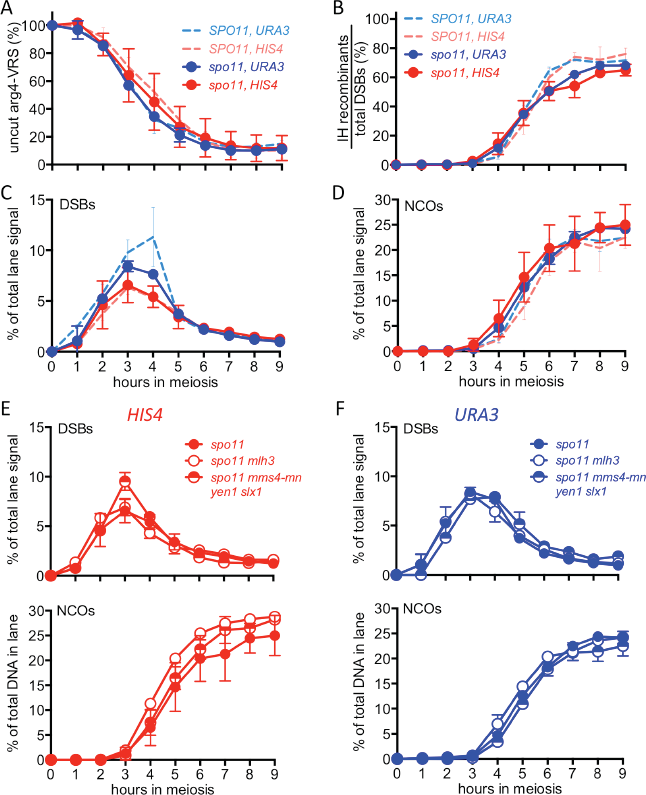
DSBs and recombinant products in *spo11* strains. (**A**) Cumulative DSB levels, expressed as loss of the VRS-containing insert, calculated as in Figure 2A. (**B**) Relative recovery of recombination products, calculated as in Figure 2— figure supplement 1B. (**C**) VDE-DSB frequencies, calculated as in Figure 2B. (**D**) NCO frequencies as in Figure 2C. In all four panels, solid lines denote data from *spo11* strains; data for wild type (dotted lines, from Figure 2 and Figure 2—figure supplement 1) are presented for comparison. (**E**) DSB (top) and NCO (bottom) frequencies in *spo11-Y135F* strains with inserts at *HIS4.* (**F**) DSB (top) and NCO (bottom) levels in *spo11-Y135F*strains with inserts at *URA3.* For all panels, values are from two independent experiments; error bars represent standard error of the mean. Representative Southern blots are in Figure 5— figure supplement 2.

**Figure 5—figure supplement 2.**
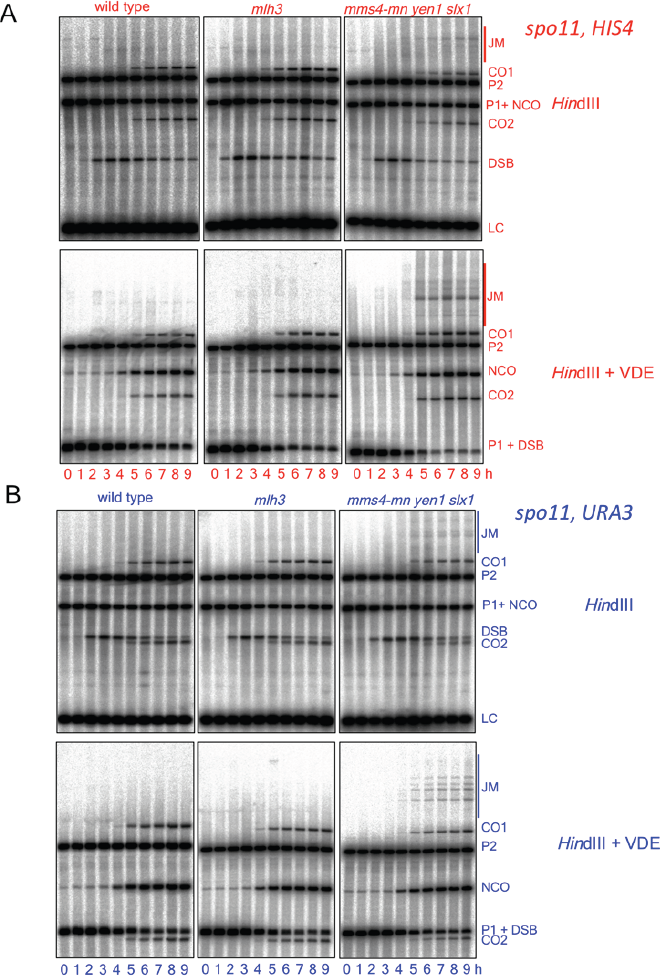
Representative Southern blots. Blots are of of *HindIII HindI*II*-VDE* digests of DNA from *spo11* strains with inserts at *HIS4* (top) and at *URA3* (bottom). Gel labels as in Figure 1; JM—joint molecule recombination intermediates.

